# Generation of raptor diversity in Europe: linking speciation with climate changes and the ability to migrate

**DOI:** 10.1101/2022.03.29.486263

**Authors:** Juan J. Negro, Eduardo Rodríguez-Rodríguez, Airam Rodríguez, Keith Bildstein

## Abstract

**Aim:** Europe should be considered as a diversity hotspot for diurnal raptors, but just during the breeding season, as it holds the higher proportion of transcontinental migratory species of any landmass, and the area becomes depleted during the winter period. This study will test the hypothesis that the high diversification of the raptor assemblage in Europe is a recent event, occurring mainly during the Quaternary, and that closely related species sharing the same trophic niches can only coexist in sympatry during the breeding period, when food availability is higher.

**Location:** Continental Europe.

**Taxon:** Diurnal birds of prey (Accipitriformes and Falconiformes).

**Methods:** A consensus molecular phylogeny for the 38 regular breeding species of raptors in Europe was obtained from BirdTree. For the same species, a trophic niche cluster dendrogram was constructed. Size and migratory strategy were introduced in the resulting phylogeny, where trophic groups were also identified.

**Results:** A total of 16 trophic groups were identified. Multispecific trophic groups tended to be composed of reciprocal sister species, while monospecific groups (only three) were composed of highly specialized species. According to time calibrated phylogenies, the speciation events took place during the glacial cycles of the Quaternary period in a majority of cases. During the non-breeding season, the smaller species in every trophic group migrate to sub-Saharan Africa, whereas larger species are either non-migratory or perform shorter migrations within Europe and/or northern Africa.

**Main conclusions:** This investigation illustrates how the rich assemblage of diurnal birds of prey in Europe, more diverse and more migratory than the North American assemblage at equivalent latitudes, has emerged recently due to the multiplication of look-alike species with similar trophic ecologies, mainly in climate refugia during cold periods. In the non-breeding season, when shared food resources are limited, smaller species migrate to Africa and alleviate competition.

## Introduction

Raptor migration is a conspicuous phenomenon in many locations globally (Bildstein, 2018). Numerous raptors, of 183 out of 313 (58%) species worldwide, abandon their breeding grounds and, often gathering in large flocks, move away at predictable times and places to reach distant areas located sometimes in different continents (Bildstein et al., 2007; Ferguson-Lees & Christie, 2001). Raptor migration has been extensively monitored to assess long-term population trends and to determine raptor flight behavior, migration corridors, as well as breeding and non-breeding grounds (Bildstein, 2006). However, much remains to be investigated concerning the causes and function of raptor migration, or why a majority of species are migratory but still others remain sedentary. Recent molecular advances have shown that there are in fact two distantly related groups of diurnal raptors, convergent in their morphology and general ecology (Jarvis et al., 2015). On one side, are the true falcons and allies, in the Order Falconiformes. On the other side, eagles, hawks, kites, harriers, and vultures constitute the Order Accipitriformes, with the New World vultures (Family Cathartidae or Order Cathartiformes, depending on the source, Del Hoyo et al., 1994; Del Hoyo, 2020) forming a sister group of all remaining Accipitriformes (Jarvis et al., 2015). According to Ericsson (2011), the early radiation of Falconiformes took place in South America, whereas Accipitriformes emerged in Africa. Today, these two orders have gained a cosmopolitan distribution and species of the two orders tend to co-occur in most environments. There are many migratory species or populations within both Falconiformes and Accipitriformes (Bildstein, 2006) in both the Nearctic and the Palearctic, implying that long-distance migratory behavior has evolved multiple times and that it is a rather plastic trait (Nagy et al., 2017).

Europe is the land mass with the highest proportion of migratory species of birds of prey in the World (Bildstein, 2006), and there is also a large absolute number of species considering its relatively small size compared to the remaining of the Palearctic and also the Nearctic (see results). Why this is so has never been explained nor explored so far. Also overlooked, even though it should be obvious glancing at any bird guide (e.g., Svensson et al., 1999) or raptor biology book (Ferguson-Lees & Christie, 2001), is the fact that there are numerous groups (mainly duets and trios) of morphologically and ecologically similar raptorial species in the European portion of the Western Palearctic. The species conforming these groups may look so similar in appearance that their field identification at a distance may be challenging (Negro, 1991; Forsman, 2016). We refer to pairs of species such as the Eurasian (*Falco tinnunculus*) and the lesser kestrels (*F. naumanni*), the lesser and greater spotted eagles (*Clanga pomarina* and *C. clanga*), or trios including, for instance, Montagu’s harrier (*Circus pygargus*), hen harrier (*C. cyaneus*) and pallid harrier (*C. macrourus*).

Here we aim to link raptor migration, interspecific competition and finally diversity and speciation in the European diurnal raptor assemblage, including both Falconiformes and Accipitriformes. Controlling for phylogeny and considering some regularities in size differences and migratory behaviour, we will test the hypothesis that the raptor community in Europe has recently experienced a diversification of breeding species, mainly during the Quaternary. Additionally, the coexistence of multiple look-alike species in the same trophic niches is only possible during the breeding season (i.e., spring and summer), when food resources are less limiting. In support of our hypothesis, that implies recent speciation events, we have gathered information on the timing of species splitting derived from time-calibrated molecular phylogenies (Fuchs et al., 2015; Mindell et al., 2018).

## Methods

### Study area and selected species

This study focuses on the 38 diurnal raptor species breeding regularly and historically in continental Europe (see Table 1). We use the term Europe as defined in the seven-continents model (i.e., Europe separated from Asia as a conglomerate of peninsulas delineated by the Mediterranean sea to the South, the Atlantic ocean to the West, the Northern sea in high latitudes, and the Ural mountains, the Caucasus and the Black and Caspian seas to the East) because it suits well the differences in migrating behaviour of birds including raptors in northern and temperate areas (e.g., Herrera, 1978; Newton & Dale, 1996; Newton & Dale, 1998, Bildstein, 2006). Using continental Europe as a biogeographical area allows us to cleanly separate sedentary, migrants within continental Europe, and sub-Saharan bird species migrating through the Mediterranean flyways (i.e., Straits of Gibraltar, Messina and Bosphorus). The Western Palearctic, a zoogeographical region widely used in bird studies (e.g., Cramp & Simmons, 1979; Perrins & Snow, 1998), and that fully encompasses Europe, introduces, however, African and Asian raptorial species that would add noise to our analyses. These species, either at the southern limit of the Western Palearctic in the African continent or at the eastern border with Asia, are all rare breeders or vagrants in the area that have never established breeding populations in continental Europe. Raptors of the Western Palearctic which are clearly African taxa include at least (Svensson et al., 1999), the lappet-faced vulture (*Torgos tracheliotos*), the dark chanting goshawk (*Melierax metabates*), the barbary falcon (*Falco pelegrinoides*), the sooty falcon (*Falco concolor*), the tawny eagle (*Aquila rapax*), and Verreaux’s eagle (*A. verreauxi*).

**Table 1.**
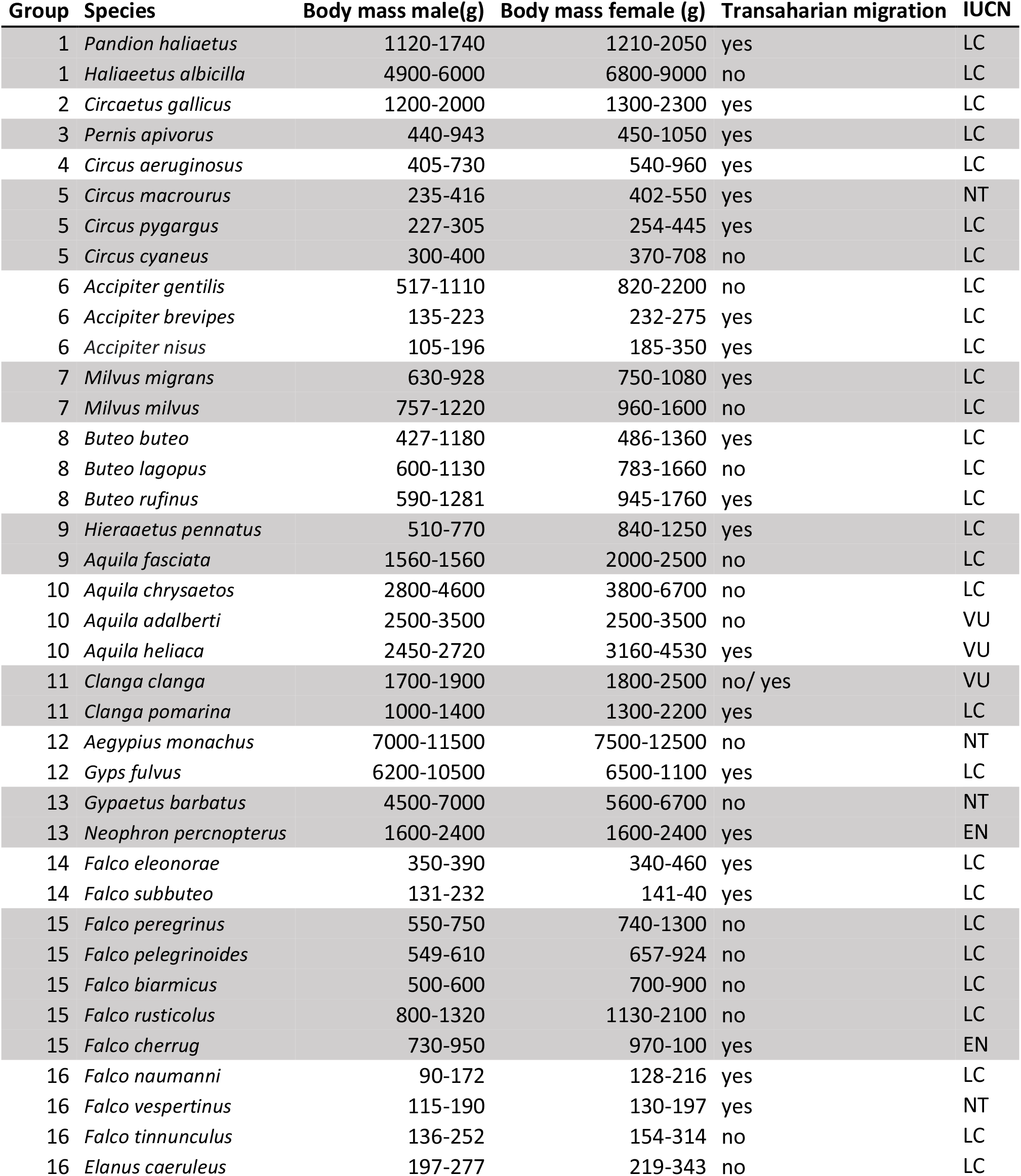
Assemblage of diurnal raptors breeding in continental Europe, ordered according to trophic niche (group, first column). Some of the groups (2 to 4) are monospecific, as they are constituted for highly niche-specific species. Range of body masses for males and females were taken from Ferguson-Lees & Christies (2001), as well as their migratory status (i.e., trans-Saharan migrant or not). IUCN conservation status is also given (IUCN 2021).

### Phylogeny and trophic niche

As a first step to determine the relationships among the different species under study, a consensus phylogeny was constructed using FigTree v 1.4.4 (Rambaut & Drummond, 2018) and 100 permutations obtained from BirdTree (Jetz et al., 2012; Jetz et al., 2014). Addressing migratory behaviour in birds requires not only the consideration of biogeographic aspects, but also a knowledge of diet specializations and prey availability (Nagy & Tokolyi, 2014). A trophic niche cluster dendrogram, based on Euclidean distances, was constructed using a dataset with data of trophic niche variables of each species, including hunting or scavenging behaviour, prey type and size (i.e., small birds, large birds, small mammals, large mammals, arthropods, herps or fishes), scavenging strategy (small pieces or big pieces) and foraging method (perch hunting, foraging in open areas, forest ambusher). These data can be consulted in Supplementary Material Appendix 1. The software utilised to construct the dendrogram was in R statistics (R Core Team, 2013). Presentation of the Figures was improved with Inkscape 1.0.1 (Harrington et al., 2020).

### Relating body mass and trans-Saharan migration

We compiled the following information of the raptor species: the minimum and maximum value of both total length and wingspan (2) from del Hoyo et al. (1994), the minimum and maximum values of length, tail, and tarsus, as well as ranges of male and female body masses (4 mass values in total) from Ferguson-Lees & Christie (2001). To estimate body mass, we averaged the minimum and maximum values for males and females. To estimate bird size, we ran a principal component analysis (PCA) on the above mentioned fourteen centered and scaled morphometric variables. The first principal component was used as a body size index. The first principal component retained 90.1% of variation (see PCA details in Table S1). The fourteen morphometric variables showed the same sign for their factor loadings and highly significant correlations to the first principal component (Figure S1). Factor loadings of the fourteen morphometric variables and importance of the components are given in Table S1. Pairwise correlations of the first principal component (body size index) and seven morphometric variables (i.e. those representing the lower ranges of each morphometric) are shown in Figure S1.

Based on results from trophic niche clustering, groupings of species sharing a similar niche were established. These groups were identified on the previously built phylogeny. Using body mass data and migratory strategy (Table 1), we established two categories: smaller raptors inside their groups and larger raptors inside their groups, and estimated the proportion of trans-continental migratory species among large and small species. Significance of the differences in frequency of migration for both groups of large and small species were tested using a contingency Table. In addition, we plotted the mass of paired migratory and resident species in the foraging and phylogenetic groups that we had previously identified (Figure 3a). By adding a diagonal line (y = x), the resulting plot graphically shows in which cases the migratory species was either larger (above the diagonal) or smaller (below the diagonal) than its resident or less-migratory counterpart. We additionally plotted the previously described body size index, because the anatomical design of migratory species is known to be slender, i.e., much longer wings, or different wing loads, compared to non-migratory species (Figure 3b).

To build the maps displayed in Figure 4, distributions ranges of raptor species were downloaded from the IUCN Red List Data (IUCN, 2021) and imported in *Qgis* (v3.20) (Qgis Development Team, 2021). We deleted passage areas for migrating species and mapped pairs and trios of species according to the analysis of trophic niche.

## Results

### Foraging groups

Based on Euclidean distances of behavioural and dietary variables, we obtained 16 trophic or ecological niche groups, including three monospecific niches (i.e., the reed-bank specialist *Circus aeruginosus,* the snake eating *Circaetus gallicus* and the himenopteran-eating *Pernis apivorus*), 7 groups of two species, 4 groups of three species, one group of 4 species, and one group of five species conformed by the bird-eating falcons (Table 1, Figure 1, Figure S2).

**Figure 1:**
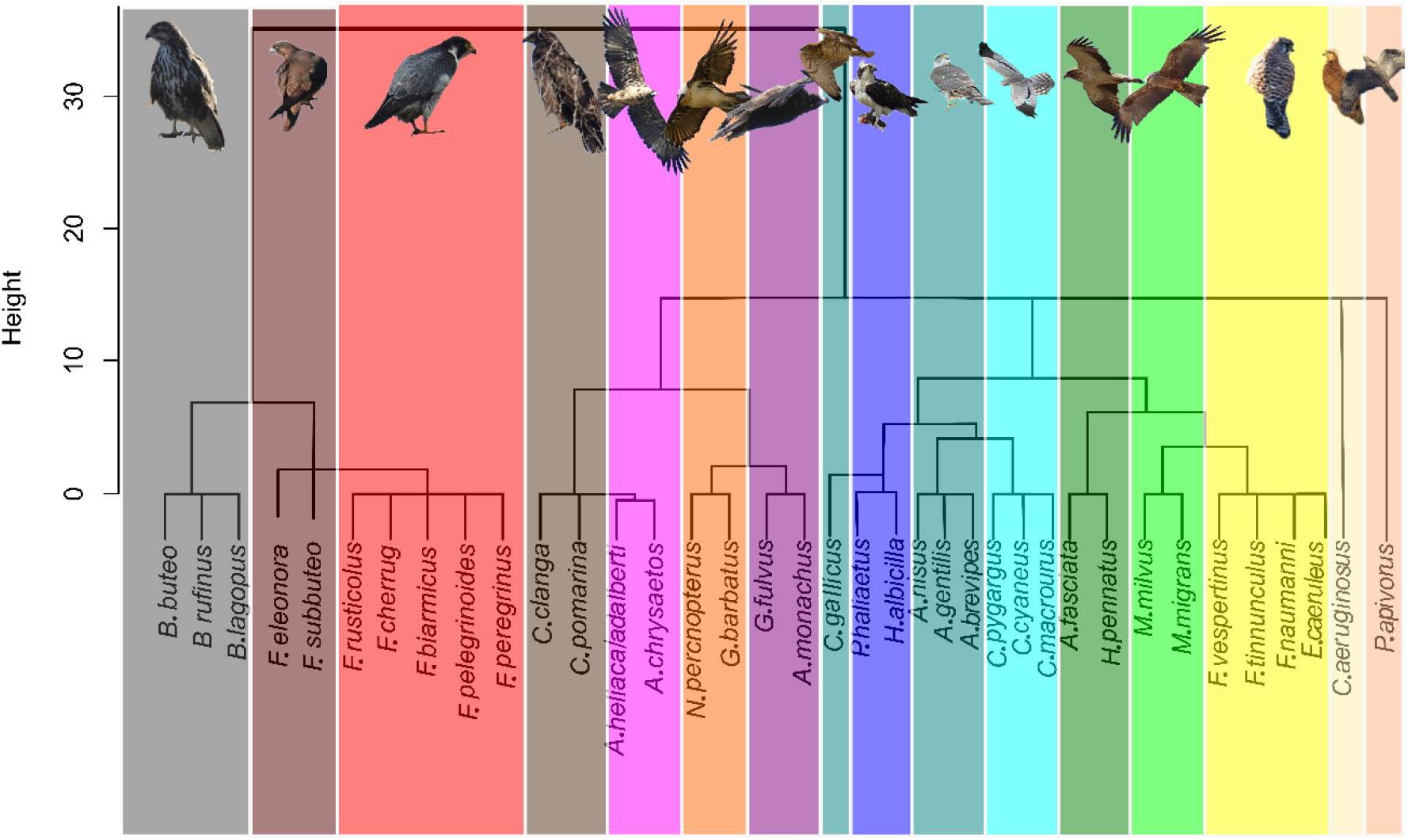
Trophic niche cluster dendrogram based on Euclidean distances for the European species of raptors. Picture credits: *Gypaetus barbatus, Accipitier nisus* and *Falco eleonorae* by Manuel Cayuela López. *Circus pygargus* by Miguel Ángel Rojas. Remaining photographs by the authors.

### Phylogenetic relationships of European raptors

The resulting phylogeny for the European raptor species matches well with the obtained trophic groups in a majority of cases, except for the *Pandion haliaetus/Haliaeetus albicilla* and *Aquila fasciata/Hieraaetus pennatus* species pairs, which are polyphyletic (Figure 2).

**Figure 2:**
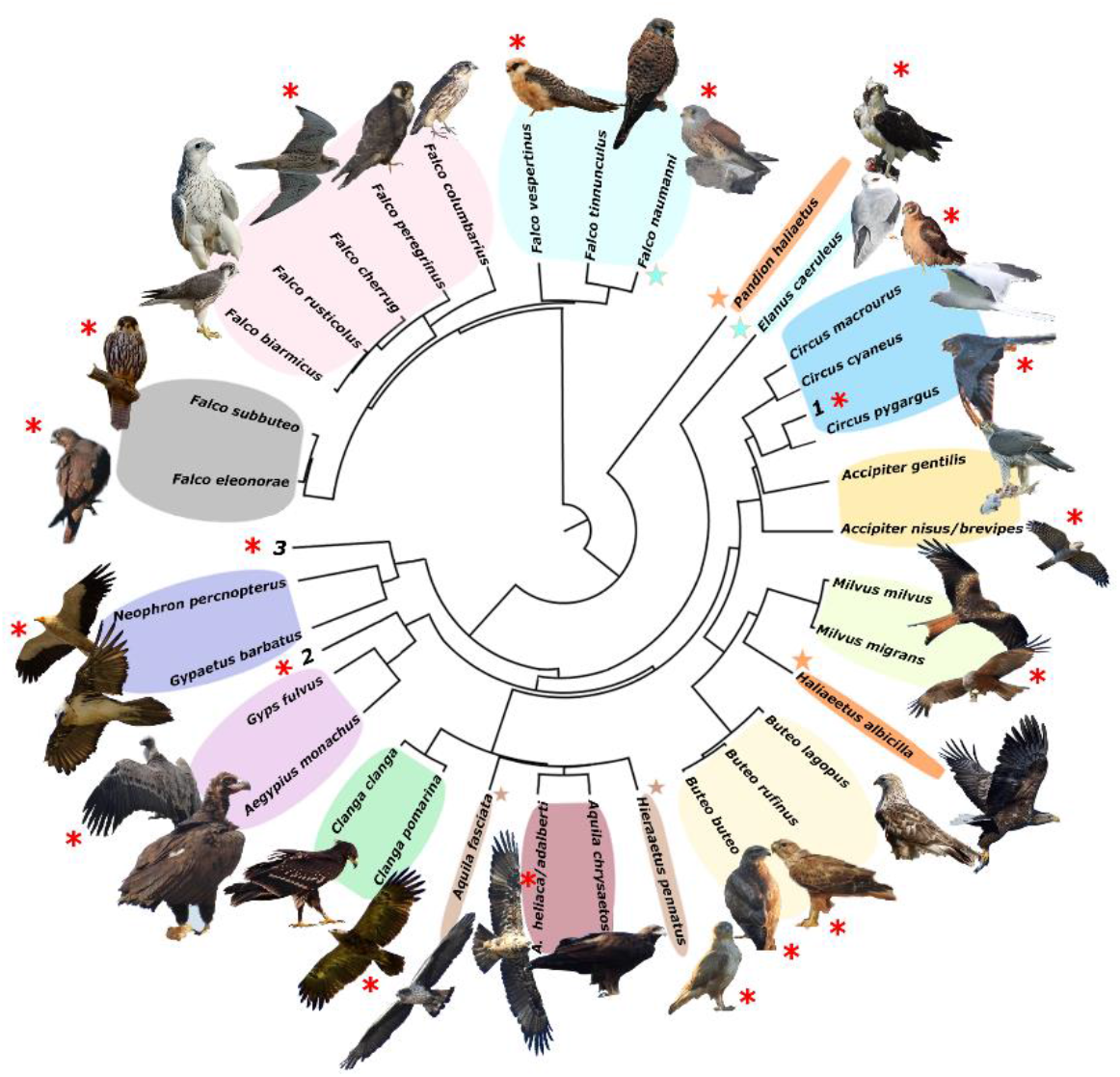
Phylogeny of included European raptors. Colour areas indicate groups of species with direct niche competition. Colour stars are used to join species not phylogenetically directly related but engage in niche competition (colour areas). Red asterisk indicated long distance migration. Number are species not include in the niche groups (1. Circus aeruginosus; 2. Circaetus gallicus; 3. Pernis apivorus). Picture credits: Clanga clanga modified from J.M. Garg (CC BY-SA 3.0). Buteo lagopus cropped from Walter Siegmund (CC BY 2.5). Haliaeetus albicilla cropped from Christoph Müller (CC BY 4.0). Falco vespertinus cropped from Martin Mecnarowski (CC BY 3.0). Falco biarmicus cropped from Derek Keats (CC BY 2.0). Falco cherrug cropped from DickDaniels (http://carolinabirds.org/) (CC BY-SA 3.0). Falco pelegrinoides cropped from Peter Wächtershäuser/ naturlichter.de (CC BY-SA 3.0). Falco eleonorae and Accipiter gentilis with permission of Manuel Cayuela López. Remaining pictures taken by the authors. License links: https://creativecommons.org/licenses/by-sa/3.0/legalcode; https://creativecommons.org/licenses/by/2.5/legalcode; https://creativecommons.org/licenses/by/4.0/legalcode; https://creativecommons.org/licenses/by/2.0/legalcode

### Implications of size and migration strategy

A majority of the smaller species inside trophic groups reach Africa during migration (81% of cases, n = 16, and only 19% do not. On the contrary, only 20% (4 out of the 20) of the larger species inside trophic groups perform a trans-Saharan migration (i.e., *Clanga clanga, Falco eleonorae, Circus macrourus* and *Buteo rufinus*). However, these “large” species, except F. eleonore, which overwinters in Madagascar, perform a much shorter migration than their smaller counterparts in their respective groups (see Figure 4). The difference in migratory behaviour among larger and smaller species is highly significant (χ^2^=11,2754, P=0.0057, see Figure 3).

**Figure 3:**
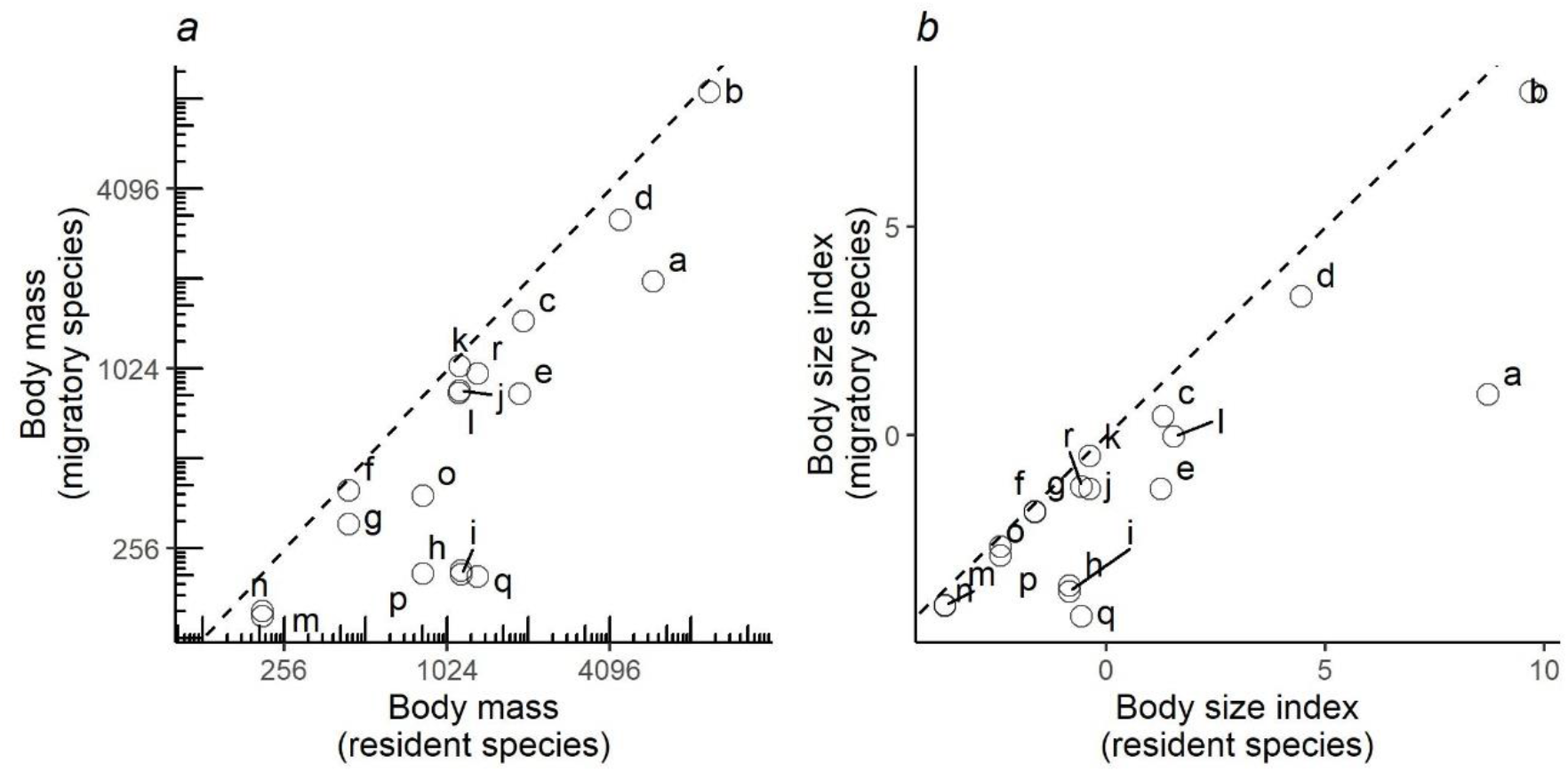
Pairwise comparisons of body mass (a) and body size (b) of raptor species derived of the foraging groups in Figure 1. Y-axis represents the migratory species in the pair, while the X-axis the non-migratory species (resident). Dashed lines indicate the diagonal (y = x). Letters refers to species pairs: a) *Gypaetus barbatus - Neophron percnopterus,* b) *Aegypius monachus* - *Gyps fulvus*, c) *Clanga clanga - Clanga pomarina*, d) *Aquila chrysaetos - Aquila heliacal*, e) *Aquila fasciata - Aquila pennatus*, f) *Circus cyaneus - Circus macrourus*, g) *Circus cyaneus - Circus pygargus,* h) *Accipiter gentilis - Accipiter brevipes,* i) *Accipiter gentilis - Accipiter nisus,* j) *Buteo rufinus - Buteo buteo,* k) *Buteo rufinus - Buteo lagopus,* l) *Milvus milvus - Milvus migrans,* m) *Falco tinnunculus - Falco naumanni,* n) *Falco tinnunculus - Falco vespertinus,* o) *Falco peregrinus - Falco eleonorae*, p) *Falco peregrinus - Falco subbuteo*, q) *Falco rusticolus - Falco columbarius*, r) *Falco rusticolus - Falco cherrug*.

When plotting the body masses for paired species derived from the foraging groups than were established above (Figure 3a), the values fall below the diagonal of the graph (x axis, mass of less migratory species, y axis, mass of more migratory species in the pair). This demonstrates that the fully or more migratory species are systematically smaller than the less migratory or resident species. Essentially the same results are obtained when instead of plotting body masses, body size indices were plotted (Figure 3b). Larger species are the less migratory ones in the species pairs. The distribution ranges for a selection of the species pairs that we considered in the above plots have been mapped in Figure 4. The less-migratory species remain within Europe (although it has to be noted that some species may have resident populations in Africa such as the peregrine falcon and the Eurasian kestrel), whereas the fully migratory species have non-overlapping breeding (in Europe) and non-breeding ranges (in Africa).

**Figure 4:**
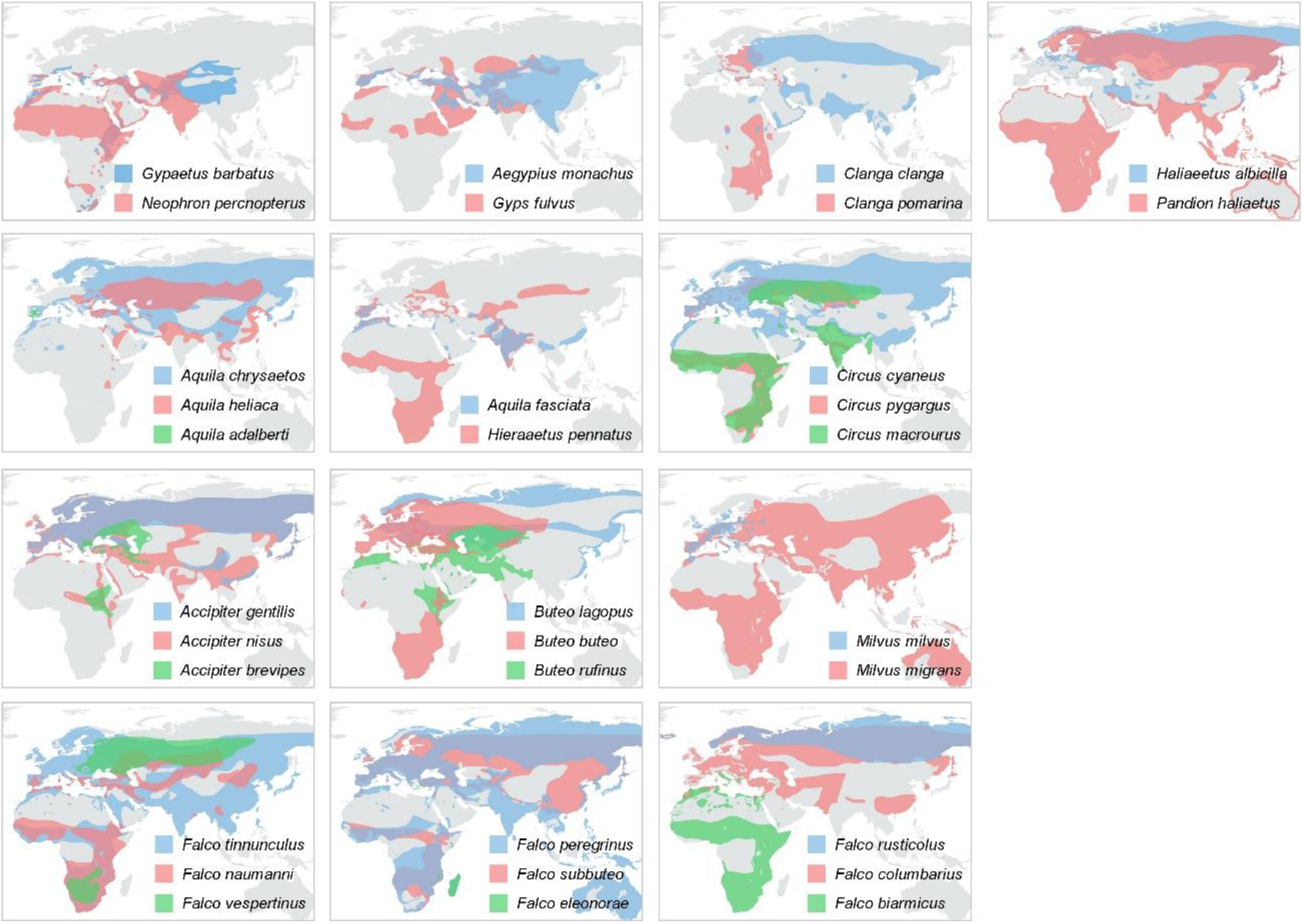
Year-round distribution maps of the raptor species with pairs or trios resulting from the trophic-niche cluster analysis. Ranges of larger in size species are displayed in blue, while ranges of the smaller and typically intercontinental migratory species are displayed in red (and green in case of trios). Distribution ranges were taken from the IUCN Red List Data (IUCN 2021).

### Timing of the splits from the common ancestor

According to the time-calibrated phylogenies of falcons (Falconiformes) on the one hand (Fuchs et al. 2015) and of all other birds of prey (eagles, hawks, vultures and kites in Accipitriformes) on the other hand (Mindell et al., 2018), six of our species groupings (both in the phylogeny and in the trophic cladogram) originated in the Quaternary (i.e., the last 2.5 million years) as sister species. These groupings are the following: 1. The red and the black kite. 2. The two spotted eagles, lesser and greater. 3. The two imperial eagles (*A. heliaca* and *A. adalberti*), that we have matched ecologically to the most distantly related *A. chrysaetos.* 4. The three European buzzards: common, rough legged and long-legged. 5. The two late-breeding falcons, dependent on the return migration of passerines for reproduction: hobby and Eleonora’s falcon. 6. The large European falcons: Lanner, Saker and Gyrfalcon. Apart from this, we may add that the two colonial and tree-nesting falcons of temperate Eurasia, i.e., the red-footed falcon in the Western Palearctic and Amur falcon in the Eastern Palearctic, which are also sister species splitting from a common ancestor whose range may have spanned the temperate forests of the whole of Eurasia.

The remaining groupings may have formed earlier in time, again according to Fuchs et al (2015) and Mindell et al. (2018): 1. the Eurasian kestrel and the lesser kestrel emerged as species in the kestrel radiation of the Pliocene, about 4-5 million years (Negro et al. 2020). 2. The harriers (hen, Montagu’s and pallid) may have radiated about 5 million years ago. 3. The hawks of the genus *Accipiter* radiated much earlier, and the common ancestor of the northern goshawk and both the European sparrowhawk and levant sparrowhawk possibly evolved 10-15 million years ago in the Miocene. 4. The two small and ventrally white eagles, Bonelli’s and booted, emerged from a common ancestor living in the Pliocene 4 million years ago. 5. The two large European vultures, griffon and cinereous, shared a common ancestor about 9 million years ago. 6. Coalescence time for Bearded and Egyptian vultures, which are sister groups to each other, is about 12 million years.

## Discussion

The diversity of species of any given taxa in a land mass typically depends on its own evolutionary history, the interaction with other species in the area, and the past and present physical patterns of the environment (e.g., Hanssen & Urban, 1992; Mönkkönen, 1994). This work was prompted by the fact that the number of breeding species of diurnal raptors in continental Europe (38 species in a land-mass of 10,180,000 km^2^) is higher than the number of breeding species of raptors in Canada and continental US combined (30 species in about 19,000,000 km^2^, including the 3 endemic New World vultures), and also by the disproportionate number of transcontinental migratory species in Europe (Bildstein 2006, see below). The latitudinal range is much wider in the American side and includes large areas below 35° North latitude in the southern states of the US that are not represented in continental Europe. However, this does not bring over a significant number of tropical species that do inhabit Mexico and Central America (the total number of raptorial species in the latter area rises to 69 species, Clark & Schmitt, 2017).

Seven species are shared between Europe and North America: five Holarctic raptor species, the gyrfalcon, the merlin, the golden eagle, the rough-legged hawk, and the northern goshawk, plus two cosmopolitan species, the peregrine falcon and the osprey, which hold breeding populations in all continents except Antarctica (Ferguson-Lees & Christie, 2001). Leaving aside these few shared species, mainly from northern or mountainous regions, there is a marked difference among Europe and Canada-US combined in the number and proportion of regular transcontinental migratory species, which is much higher in Europe (22 species crossing the Mediterranean Sea to Africa in the non-breeding season) than in the above mentioned area of North America, where only five species make the bulk of the migrants moving down to South America (i.e., *Ictinia mississippiensis, Buteo swainsoni, B. platypterus, Cathartes aura* and *Pandion haliaetus*) (Bildstein, 2006).

Our results show that there is a duplicity, or even multiplicity of species in the same trophic and ecological niches in Europe. In many cases the species involved are so similar that identification at a distance may be difficult (Forsman, 2016). This similarity may be due to shared ancestry in congeneric species, such as in the duet conformed by *C. pomarina* and *C. clanga,* or in the case of *Milvus milvus* and *M. migrans* (Mindell et al., 2018), but it may be due to convergence in more distantly related species such as *A. fasciata* and *H. pennatus*. Duplicated species are not apparent in North America, where the two more diverse genera (*Buteo* and *Falco*), have rather distinct species in terms of size and ecology. That Europe has been a recent hotspot for raptor speciation is also supported by the pattern of endemism (Weir and Schluter, 2004) in the group: six species are only or mainly distributed in Europe (i.e., red kite, honey buzzard, lesser spotted eagle, Spanish imperial eagle, Montagu’s harrier and Eleonora’s falcon). Five other species breed in Europe and more into the East, but may have a Western Palearctic origin: lesser kestrel (with the older fossil found in the Iberian Peninsula, Negro et al., 2020), red-footed falcon (sister species of the Amur falcon breeding in eastern Asia, as commented above), common buzzard (the nominal subspecies is sedentary across Europe, and only the Eastern subspecies *B. b. vulpinus* migrates to Africa), and finally the levant sparrowhawk, and the western marsh harrier.

Although our work does not focus on the evolution of raptor migration, our results shed light on the biogeographical origin of at least some of the migratory species. A majority of the duplicated species would adhere to the “northern origin” hypothesis (Bell, 2000; Bruderer and Selewski, 2008, Nagy et al., 2014), according to which harsh winter conditions would impel northern species to move to Africa in the non-breeding season. Some species in the system, such as the short-toed eagle, the Egyptian vulture and the bearded vulture, and two small eagles, Bonelli’s and booted eagles, may have an afro-tropical origin, as they belong to phylogenetic groups that are highly diverse in Africa (Mindell et al., 2018). None-the-less, the migratory species among them may have acquired the migratory behaviour after colonizing Europe and competing with other species there.

Taken together, our results support the notion that the current raptor assemblage in Europe experienced a recent enrichment, and may be considered a hotspot accounting for its area and geographic location, and particularly by comparison with the smaller raptor community in much larger North America. This does not mirror the situation for the whole of the avifaunas, as there are about 556 bird species in Europe (Keller et al. 2021), compared to 740 in North America to the north of Mexico (Cornell lab of Ornithology, https://www.notesfromtheroad.com/roam/how-many-birds-north-america.html). The North American avifauna contains a plethora of insectivorous passerines with a probable tropical or subtropical origin (i.e., Parulidae, Turannidae, Vireonidae, Icteridae) which exploit the insect-rich summer of North America and migrate to the south in the winter (Snow 1978). The European migratory songbirds are fewer and belong to families (i.e., Sylviidae, Turdidae, Muscicapidae and Motacillidae) whose evolutionary centres are Palearctic but not African (Snow 1978). As a result, the assemblages of forest passerine birds of the Nearctic region are more diverse than those of the Western Palearctic, thus including Europe (Mönkkönen 1994). Other taxa follow the same pattern: Western North America contains, for instance, 41% more mammal species than Europe (Mönkkönen & Viro, 1997).

One mechanism that may explain the generation of raptor diversity in Europe is the existence of climate-driven vicariant events during the glacial cycles of the Pleistocene (Voelker 2010), and before during the Pliocene or earlier in the Miocene. It has been suggested, for instance, that the genus *Falco* radiated during the Miocene several million years ago, coincidental with a period of increased aridity favouring grasslands (Zachos et al., 2001, Fuchs et al. 2015). Glacial cycles have been invoked as drivers of speciation for numerous bird taxa in both the Old and the New World: fragmentation of populations, isolation in refugia during glacial times and subsequent reunion without admixture is a common model for explaining the origin of closely-related bird species (Bermingham et al., 1992 Weir and Schluter, 2004, Lovette, 2005). Such a model was in fact invoked to explain the emergence as separate species of the Eastern Imperial Eagle and the Spanish imperial eagle (Ferrer and Negro, 2004). The latter is the species in the genus *Aquila* with the smallest distribution range, as it is endemic to the Iberian peninsula.

Lacking enough dated fossils, the recent avian speciation during the Pleistocene glacial cycles has remained controversial for some groups (Lovette, 2005; Zink et al., 2004; Wang et al., 2018). However, recent advances in molecular techniques, such as whole-genome sequencing, clearly support that many bird species suffered dramatic expansions and contractions during the Quaternary coincidental with climate changes (Nadachowska-Bryzska et al., 2015). These population changes may have prompted speciation.

The generation of sister species of birds of prey may be explained by vicariance events. The difference among Europe and North America possibly has to do with the different effects of the glaciation in the two land masses, and the alignment of the major mountain barriers that may have acted as geographical barriers, with a north-south component in the Nearctic and east-west in the Palearctic (Albach et al., 2006). In North America, numerous pairs of sister bird species tend to have east-west distributions (e.g., Lovette, 2005). In Europe, the existence of several southern peninsulas and islands offered different refugia and thus more possibilities for population fragmentation and isolation (Covas & Blondel, 1998). After ice cover retreated during the interglacials, isolated populations might have expanded and established contact but, in some cases, without genetic admixture. However, closely related species of predators coexisting in sympatry would compete dearly in the harsh winter season. Competition would be resolved by the less competitive species moving away and thus migrating to a different continent. This may also be seen as several concurrent cases of character displacement, with competing species diverging in size precisely to diminish potential competition (Dayan & Simberloff, 1998). Also of interest may the fact that the assemblage of European carnivores (Mammalia) is much smaller, with 22 species than the North American assemblage, with 45 species (Burgin et al., 2020). Lessened potential competition with mammalian carnivores may have facilitated the diversification of European breeding raptors. None-the-less, competition with other predators alone is not the only possible explanation, as Bergmann’s rule (Bergmann 1847), for instance, would also predict larger bird species having a thermoregulatory advantage compared to smaller ones during the cooler non-breeding season (Salewski & Watt, 2017). As additional support to our hypothesis, the three New World vultures of North America conform to the size and migration “rules”, with the largest species, the California Condor, being sedentary, and the much smaller species turkey and black vultures being migratory (Ferguson-Lees & Christie 2001).

An extraordinary case of rapid increase of size in the absence of competitors, therefore linking body size to competition in the raptor group, is provided by two extinct birds of prey in New Zealand (Knapp et al., 2019): the largest known eagle in the world, Haast’s eagle *Hieraaetus moorei,* and the largest known harrier in the world, Eyles’ harrier *Circus teauteensis.* Both derived from Australian vagrants, the much smaller *Hieraaetus morphnoides* and *Circus assimilis*, respectively. These two concurrent cases of island gigantism occurred in the early Pleistocene when the clearing of closed forests due to climate changes allowed for successful colonization of Australian raptor immigrants hunting in open woodlands and grasslands (Knapp et al., 2019). The fact that body size is in turn related to migration propensity and ability is also supported at the intraspecific level. Raptors tend to present reversed sexual dimorphism, as females are larger than males (Newton, 1979). Numerous raptor species show different migration timings and non-breeding distributions, although it is true that there is variation in which sex migrates first (both out-bound and in-bound) or overwinters in more northerly locations (see Bildstein, 2006).

Finally, we call attention to the fact that most raptorial species in Europe have vulnerable populations (Burfield, 2008) and some, as with the Spanish imperial eagle and the bearded vulture, were on the brink of extinction in the late 20th century (Bustamante, 1998; Martínez-Cruz et al., 2004). The system we have described here seems to be unique to Europe and has resulted in a hotspot for raptors, many of which are migratory and face conservation problems both inside and outside Europe. The current scenario of climate change adds uncertainty to population persistence (Rodríguez-Rodríguez et al., 2020), while some trans-Saharan migratory species are already changing their behaviour and becoming non-migratory in southern Europe. This includes at least the Egyptian vulture (Di Vittorio et al. 2016), lesser kestrel (Negro et al. 1991), booted eagle (Mellone et al., 2013) and short-toed eagle (Martínez & Sánchez-Zapata 1999). In addition, African species are entering Europe from the south, as it possibly happened with the black-shouldered kite in historical times (Balbontín et al., 2008) and it is just starting with the Ruppell’s vulture (Ramírez et al., 2011). If interspecific competition was central in explaining diversity, distribution and migratory behaviour in the European raptor community, as our results suggest, the rapid contemporary changes in climate and bird’s ecology will favour some species at the expense of others, generating new conservation challenges.

## Supporting information

Supplementary matterial

## Acknowledgements

We thank M.A. Rojas, M.Cayuela, J.M. Irastorza, J.M. Sayago and A. Kovacs for sharing pictures included in Figures 1 and 2, and Figure S2. We also thank all the photographers who have shared pictures under Creative Commons licenses. This study received no funding.

## Biosketch

**Juan José Negro** has been investigating raptor ecology and behavior for more than three decades. He was Director of Doñana Biological Station-CSIC in Spain, and currently holds a position as Research Professor in the Spanish National Research Council.

## Authors’ contributions

JJN conceived the hypothesis, wrote the first draft of the manuscript and revised the last versions. EJJRR analyzed the trophic niche data and generated trophic niche and phylogeny of European raptors, contributing also to the method and results sections. AR performed the body mass analyses in relation with migration, prepared distribution maps and contributed with manuscript writing and editing. KB provided insights, and numerous references. All authors helped to draft the paper and edited the text.

