## Supplementary matterial for "Generation of raptor diversity in Europe: linking speciation with climate changes and the ability to migrate"

^1^Estación Biológica de Doñana-CSIC, Avda. Americo Vespucio 26. 41092. Sevilla. Spain.

^2^Department of Integrated Sciences, Faculty of Experimental Sciences, University of Huelva, Huelva, Spain.^.^

^3^Grupo de Ornitología e Historia Natural de las islas Canarias (GOHNIC), C/La Malecita s/n, Buenavista del Norte, Canary Islands, Spain.

^4^Terrestrial Ecology Group (TEG-UAM), Department of Ecology, Universidad Autónoma de Madrid, Madrid, Spain.

^5^Centro de Investigación en Biodiversidad y Cambio Global (CIBC-UAM), Universidad Autónoma de Madrid, Madrid, Spain.

^6^116 Village Drive, Blandon, PA 19510, USA

**Content**

**Table S1.** Factor loadings and importance of the components of the principal components analysis (PCA) on fourteen morphometric variables.

**Table S2.** Trophic niche variables used in the analysis of trophic niche.

**Figure S1.** Pairwise correlations of the first principal component (body size index) and seven morphometric variables (i.e. those representing the lower ranges of each morphometric). Black numbers indicate Pearson correlation coefficients and red stars indicate if the correlation is highly significant (*** < 0.001).

**Figure S2.** European raptors grouped according to both trophic niche and phylogenetic relationships to show morphological and plumage similarities.

**Table S1.**

|  | PC1 | PC2 | PC3 | PC4 | PC5 |
| --- | --- | --- | --- | --- | --- |
| *Factor loadings* | |  |  |  |  |
| Size_HBW1^a^ | -0.276 | 0.136 | -0.029 | 0.311 | -0.084 |
| Size_HBW2 ^a^ | -0.275 | 0.088 | -0.096 | 0.190 | 0.499 |
| Wingspan_HBW1 ^a^ | -0.273 | 0.103 | 0.425 | -0.061 | -0.156 |
| Wingspan_HBW2 ^a^ | -0.261 | 0.032 | 0.721 | -0.413 | 0.089 |
| Length1 ^b^ | -0.278 | 0.050 | -0.024 | 0.241 | -0.181 |
| Length2 ^b^ | -0.277 | 0.087 | -0.117 | 0.084 | 0.240 |
| Wingspan1 ^b^ | -0.276 | 0.067 | 0.119 | 0.397 | -0.196 |
| Wingspan2 ^b^ | -0.276 | 0.076 | 0.090 | 0.341 | 0.216 |
| Tail1 ^b^ | -0.253 | 0.429 | -0.249 | -0.215 | -0.465 |
| Tail2 ^b^ | -0.249 | 0.437 | -0.345 | -0.477 | 0.202 |
| Weight_M1 ^b^ | -0.263 | -0.374 | -0.107 | -0.080 | -0.158 |
| Weight_M2 ^b^ | -0.260 | -0.381 | -0.155 | -0.096 | -0.419 |
| Weight_F1 ^b^ | -0.263 | -0.349 | -0.146 | -0.233 | 0.239 |
| Weight_F2 ^b^ | -0.260 | -0.401 | -0.123 | -0.103 | 0.147 |
| *Importance of components* | |  |  |  |  |
| Standard deviation | 3.563 | 0.891 | 0.472 | 0.314 | 0.252 |
| Proportion of variance | 0.907 | 0.057 | 0.016 | 0.007 | 0.005 |
| Cumulative proportion | 0.907 | 0.964 | 0.979 | 0.986 | 0.991 |

**Table S1.** Factor loadings and importance of the components of the principal components analysis (PCA) on 14 morphometric variables. ^a^ and ^b^ taken from Del Hoyo et al. (1994) and Ferguson-Lees & Christie (2001).

**Table S2**

| **Species** | **Predator** | **Scavenger** | **Prey fish** | **Prey small birds** | **Prey big birds** | **Prey small mammals** | **Prey big mammals** | **Prey insects** | **Prey herps** | **Scavenging type** | **Foraging method** |
| --- | --- | --- | --- | --- | --- | --- | --- | --- | --- | --- | --- |
| ***Pandion haliaetus*** | yes | no | yes | no | no | no | no | no | no | no | open foraging in water |
| ***Haliaeetus albicilla*** | yes | no | yes | no | yes | no | yes | no | no | no | open foraging in water |
| ***Elanus caeruleus*** | yes | no | no | yes | no | yes | no | yes | yes | no | open foraging |
| ***Circaetus gallicus*** | yes | no | no | no | no | no | no | no | yes | no | perch-hunting |
| ***Pernis apivorus*** | yes | no | no | no | no | no | no | yes | no | no | bee hunting |
| ***Circus aeruginosus*** | yes | no | yes | yes | yes | yes | no | no | no | small pieces | open foraging over wetlans |
| ***Circus pygargus*** | yes | no | no | yes | no | yes | no | yes | yes | no | open foraging close to ground |
| ***Circus cyaneus*** | yes | no | no | yes | no | yes | no | yes | yes | no | open foraging close to ground |
| ***Circus macrourus*** | yes | no | no | yes | no | yes | no | yes | yes | no | open foraging close to ground |
| ***Accipiter gentilis*** | yes | no | no | yes | yes | yes | no | no | no | no | forest ambush |
| ***Accipiter brevipes*** | yes | no | no | yes | yes | yes | no | no | no | no | forest ambush |
| ***Accipiter nisus*** | yes | no | no | yes | yes | yes | no | no | no | no | forest ambush |
| ***Milvus migrans*** | yes | yes | no | yes | no | yes | no | yes | yes | small pieces | open foraging |
| ***Milvus milvus*** | yes | yes | no | yes | no | yes | no | yes | yes | small pieces | open foraging |
| ***Buteo buteo*** | yes | yes | no | yes | yes | yes | no | no | yes | small pieces | perch-hunting |
| ***Buteo lagopus*** | yes | yes | no | yes | yes | yes | no | no | yes | small pieces | perch-hunting |
| ***Buteo rufinus*** | yes | yes | no | yes | yes | yes | no | no | yes | small pieces | perch-hunting |
| ***Hieraaetus pennatus*** | yes | no | no | no | yes | no | yes | no | yes | no | open foraging |
| ***Aquila fasciata*** | yes | no | no | no | yes | no | yes | no | yes | no | open foraging |
| ***Aquila chrysaetos*** | yes | yes | no | no | yes | no | yes | no | yes | Big pieces | open foraging |
| ***Aquila adalberti*** | yes | yes | no | no | yes | no | yes | no | yes | Big pieces | open foraging |
| ***Aquila heliaca*** | yes | yes | no | no | yes | no | yes | no | yes | Big pieces | open foraging |
| ***Clanga clanga*** | yes | yes | no | no | yes | no | yes | no | yes | Big pieces | open foraging |
| ***Clanga pomarina*** | yes | yes | no | no | yes | no | yes | no | yes | Big pieces | open foraging |
| ***Aegypius monachus*** | no | yes | no | no | no | no | no | no | no | Big pieces | open foraging |
| ***Gyps fulvus*** | no | yes | no | no | no | no | no | no | no | Big pieces | open foraging |
| ***Gypaetus barbatus*** | no | yes | no | no | no | no | no | no | no | small pieces | open foraging |
| ***Neophron percnopterus*** | no | yes | no | no | no | no | no | no | no | small pieces | open foraging |
| ***Falco naumanni*** | yes | no | no | yes | no | yes | no | yes | yes | no | open foraging |
| ***Falco vespertinus*** | yes | no | no | yes | no | yes | no | yes | yes | no | open foraging |
| ***Falco tinnunculus*** | yes | no | no | yes | no | yes | no | yes | yes | no | open foraging |
| ***Falco eleonorae*** | yes | no | no | yes | yes | yes | no | no | yes | no | perch-hunting |
| ***Falco subbuteo*** | yes | no | no | yes | yes | yes | no | no | yes | no | perch-hunting |
| ***Falco peregrinus*** | yes | no | no | yes | yes | yes | yes | no | yes | no | perch-hunting |
| ***Falco columbarius*** | yes | no | no | yes | yes | yes | yes | no | yes | no | perch-hunting |
| ***Falco biarmicus*** | yes | no | no | yes | yes | yes | yes | no | yes | no | perch-hunting |
| ***Falco rusticolus*** | yes | no | no | yes | yes | yes | yes | no | yes | no | perch-hunting |
| ***Falco cherrug*** | yes | no | no | yes | yes | yes | yes | no | yes | no | perch-hunting |

**Table S2.** Behavioural and dietary data used to generate trophic niche groups.

**Figure S1**

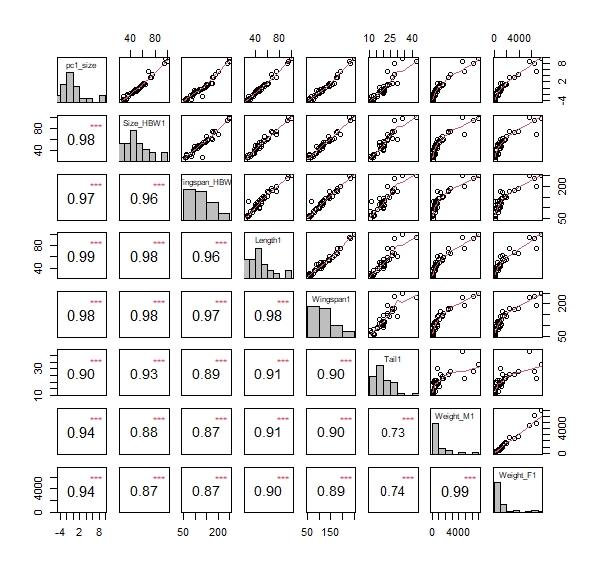

**Figure S1.** Pairwise correlations of the first principal component (body size index) and seven morphometric variables (i.e. those representing the lower ranges of each morphometric). Black numbers indicate Pearson correlation coefficients and red stars indicate if the correlation is highly significant (*** < 0.001).

**Figure S2.**

**Group 1**: *Haliaeetus albicilla* (left) by C. Müller (Creative Commons). *Pandion haliaetus* (right) by J.A. Irastorza.
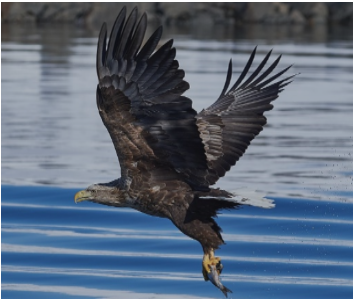

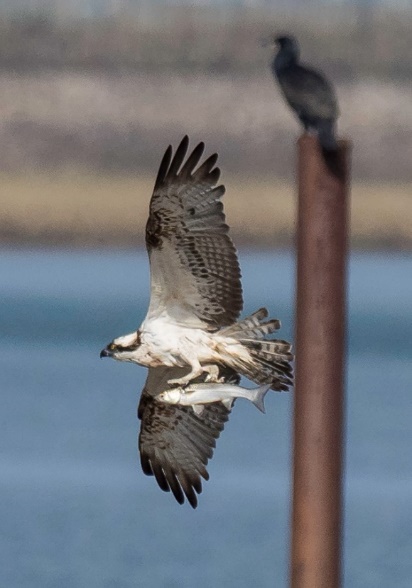

**Group 2**: *Circus macrourus* (left) by M. Rojas. *C. pygargus* (center) by J.J. Negro. *C. cyaneus* (right) by B. Rodríguez.
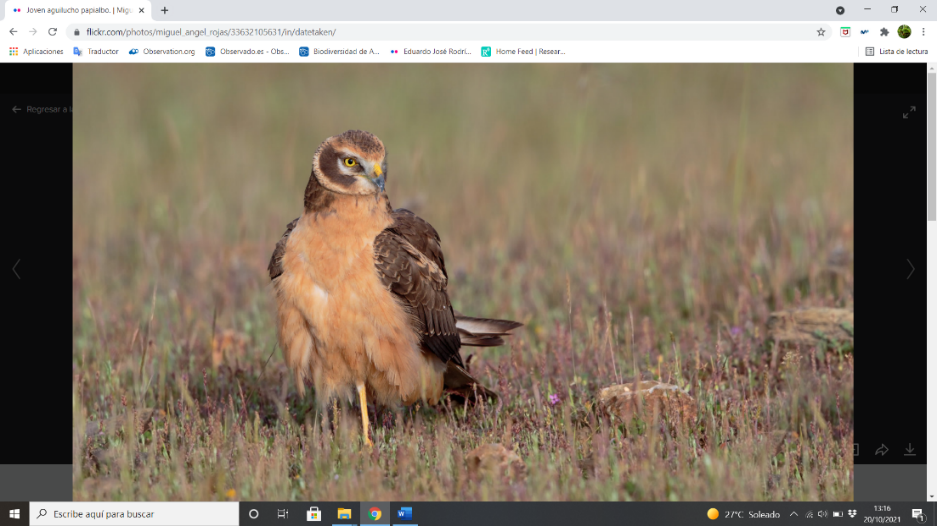

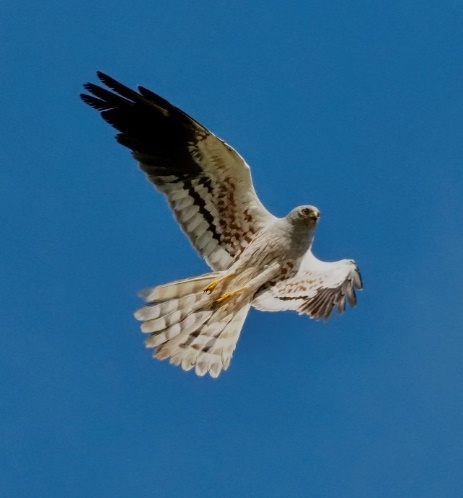

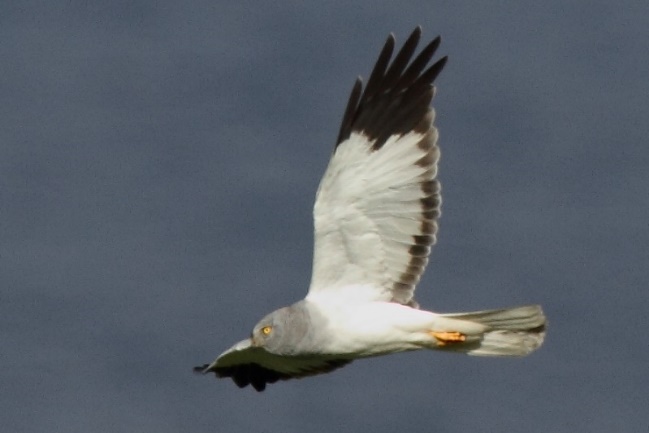

**Group 3**: *Accipiter gentilis* (left) and *A. nisus* (center) by M. Cayuela*. A. brevipes* (right) by V. Y. Arkhipov (Creative Commons).

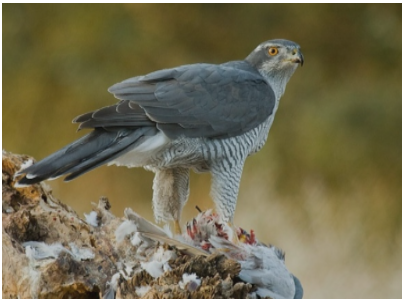

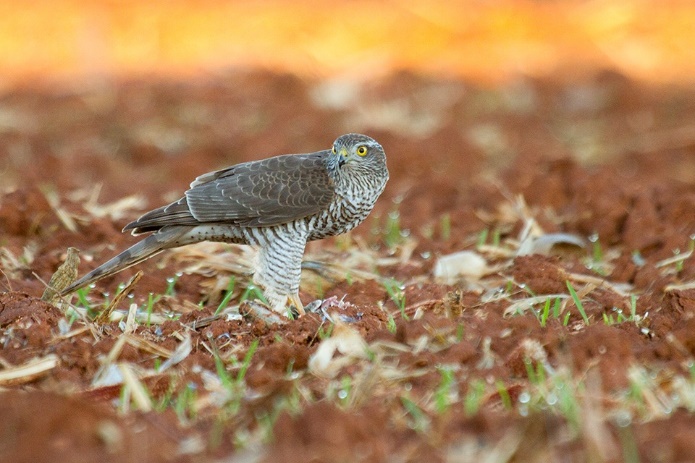

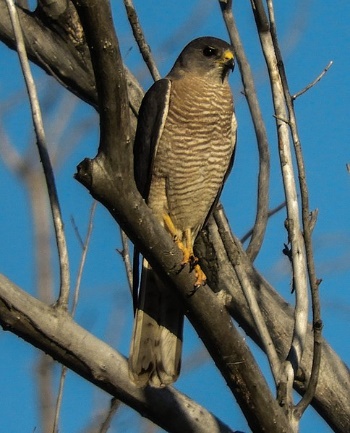

**Group 4**: *Milvus milvus* (left) and *M. migrans* by J.J. Negro.

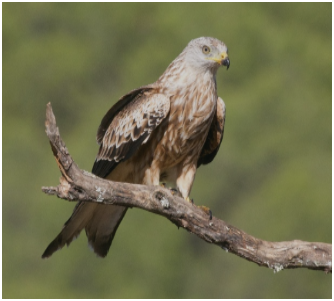

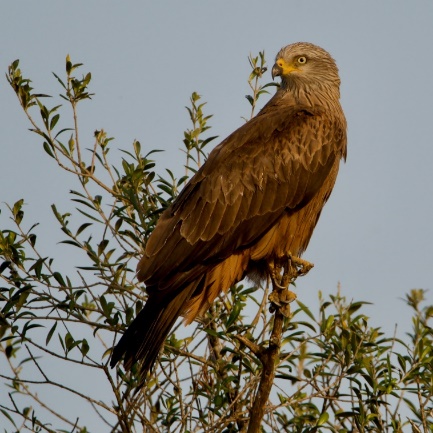

**Group 5**: *Buteo lagopus* (left) by M. Szczepanek (Creative Commons). *B. buteo* (center) by E.J. Rodríguez-Rodríguez. B. *rufinus* (right) by K. Koshy (Creative Commons).

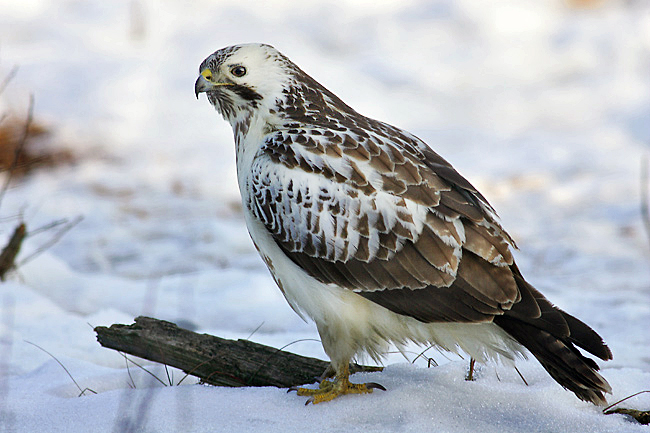

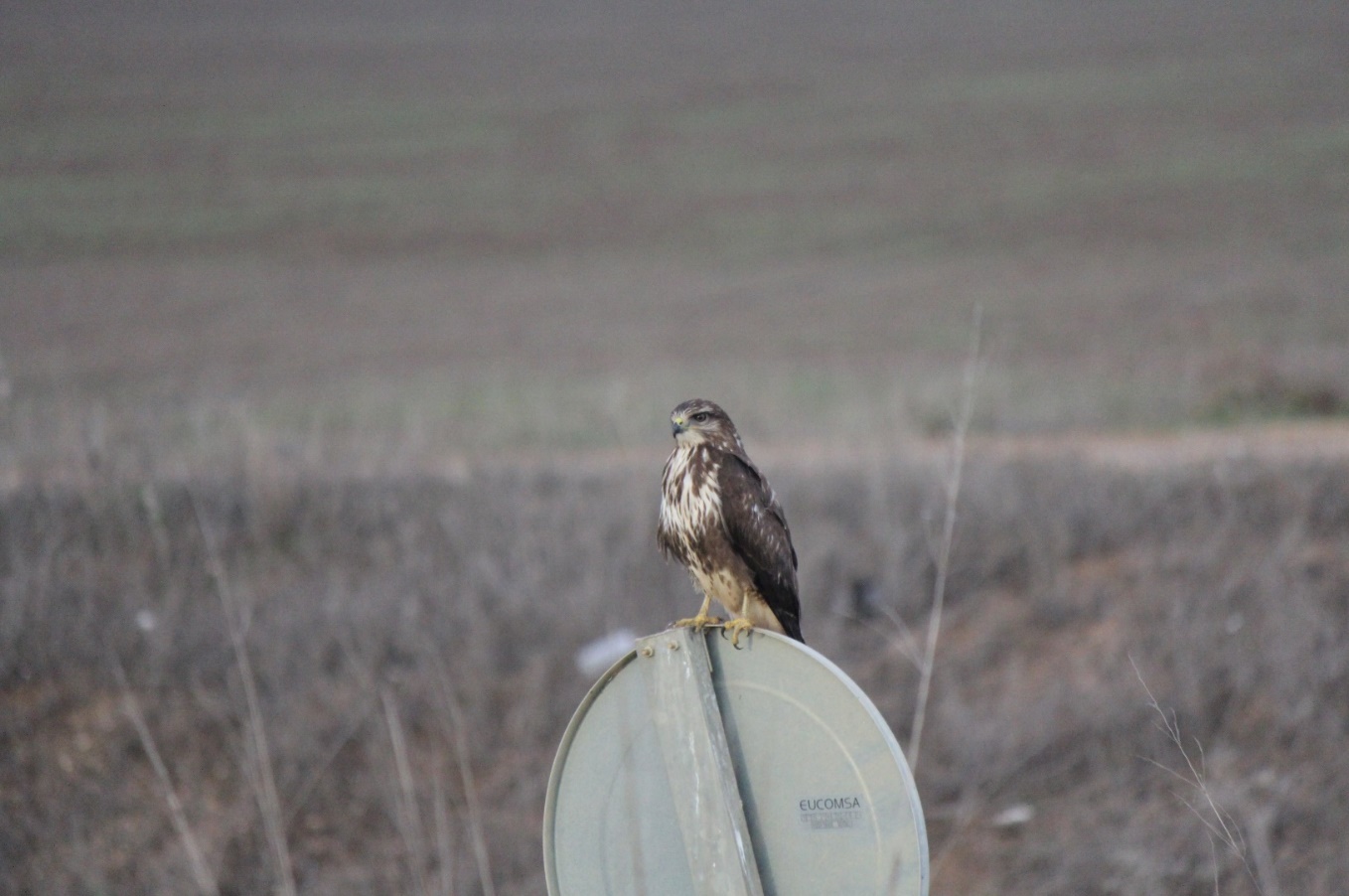

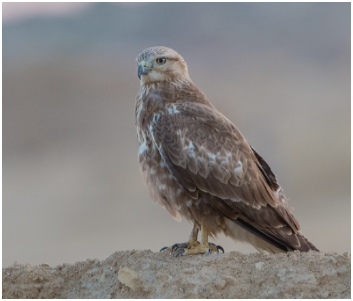

**Group 6**: *Aquila fasciata* (left) by J.A. Irastorza. *Hieraaetus pennatus* (right) by J.J. Negro.

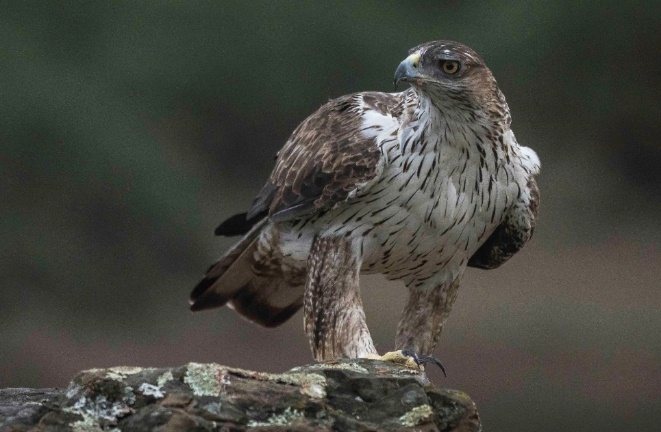

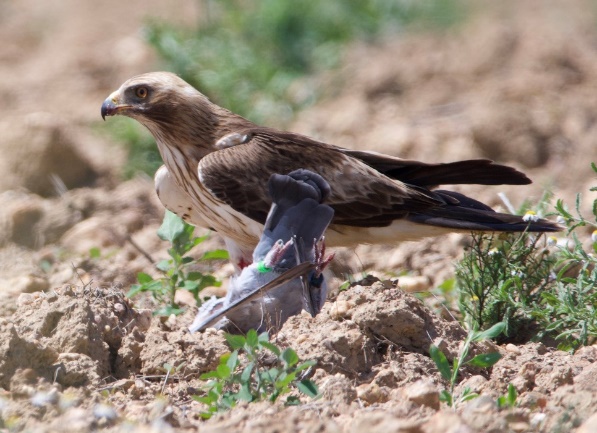

**Group 7**: *Aquila chrysaetos* (left) and *A. adalberti* (center) by J.J.Negro. *A. heliaca* (right) by A. Kovacs.

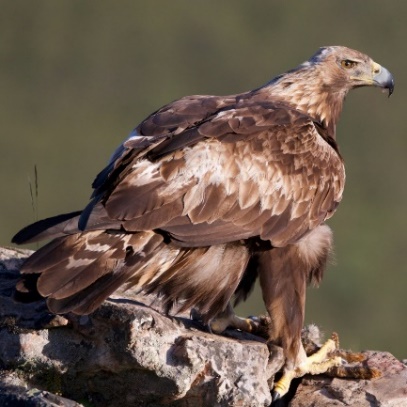

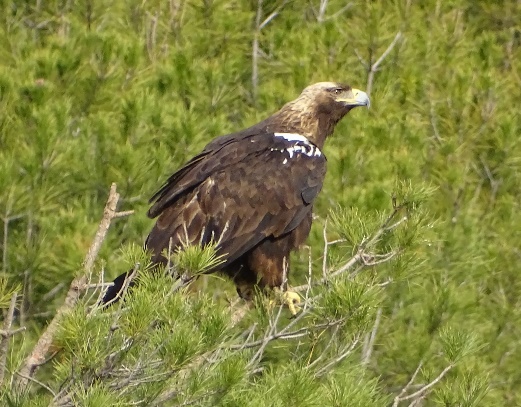

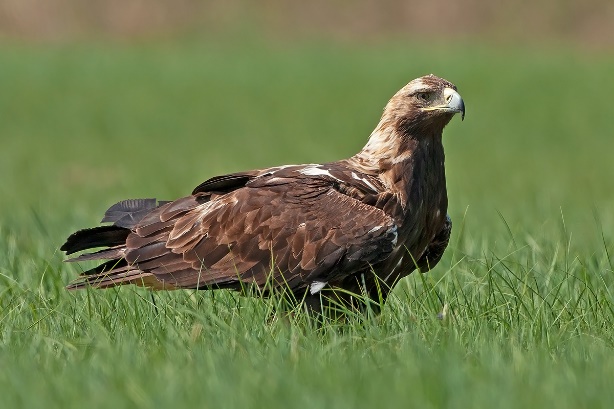

**Group 8**: *Clanga clanga* (left) by A. Kovacs. *C. pomarina* (right) by W. Avnerunder (Creative Commons). Juvenile plumage (both).

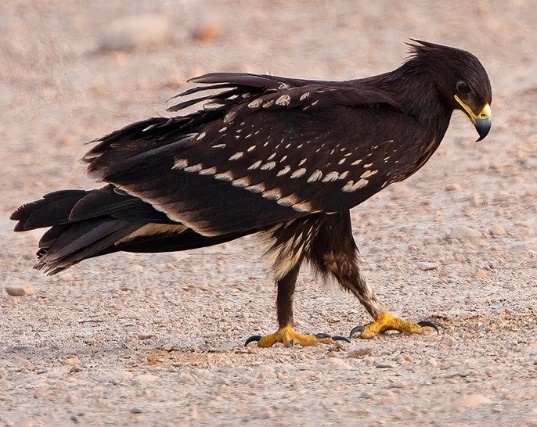

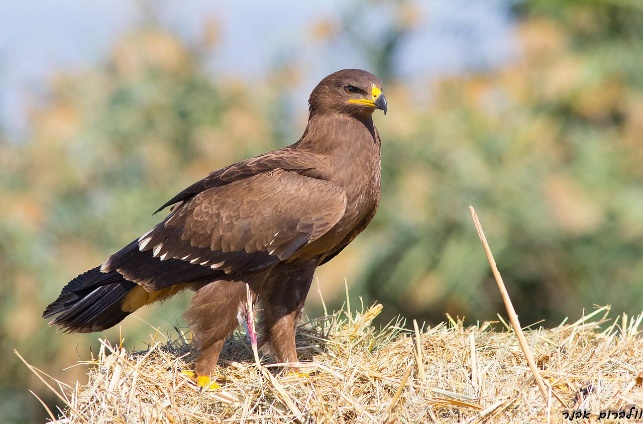

**Group 9**: *Gyps fulvus* (left and center) *and Aegypius monachus* (right) by J.J. Negro.

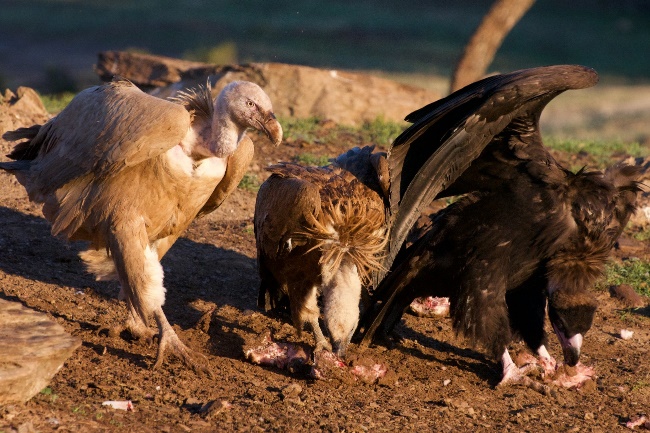

**Group 10**: *Gypaetus barbatus* (left) and *Neophron percnopterus* (right) by J.J. Negro.

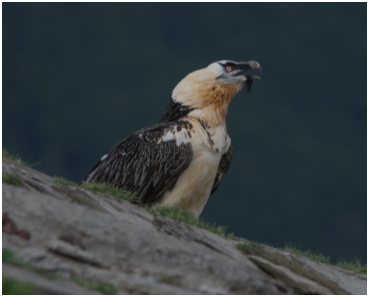

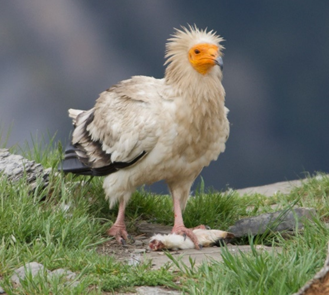

**Group 11**: *Falco eleonorae* (left) by B. Rodríguez. *F. subbuteo* (right) by J.J. Negro.

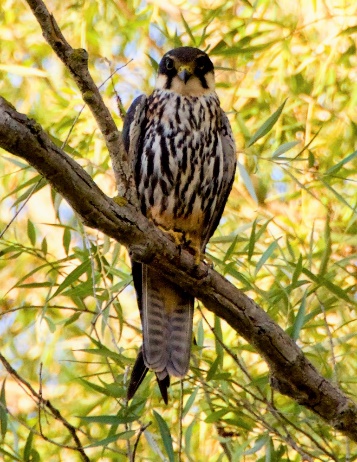

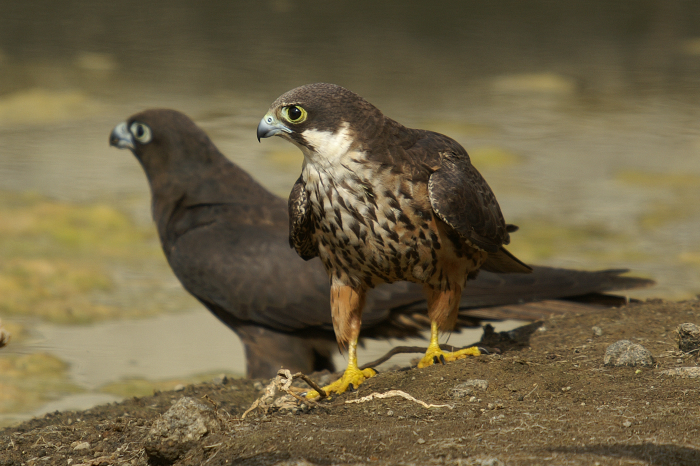

**Group 12**: *Falco rusticolus* (left) by NorthernLight (Creative Commons). *F. cherrug* (center) by B. Rodríguez. *F. peregrinus* (right) by J.M. Sayago. *F. biarmicus* (right second row) by D. Keats (Creative Commons). *F. columbarius* by G. Smith (Creative Commons).

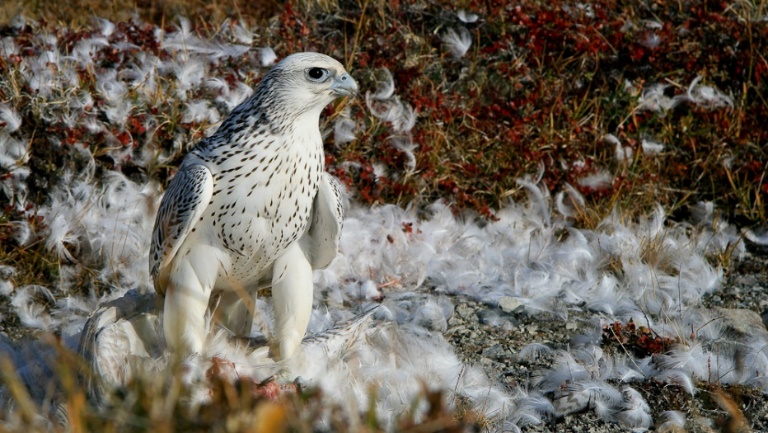

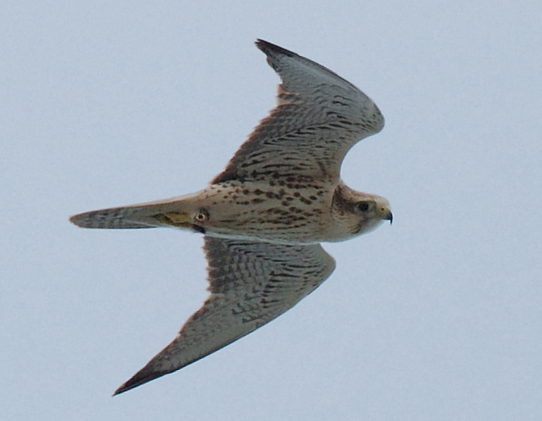

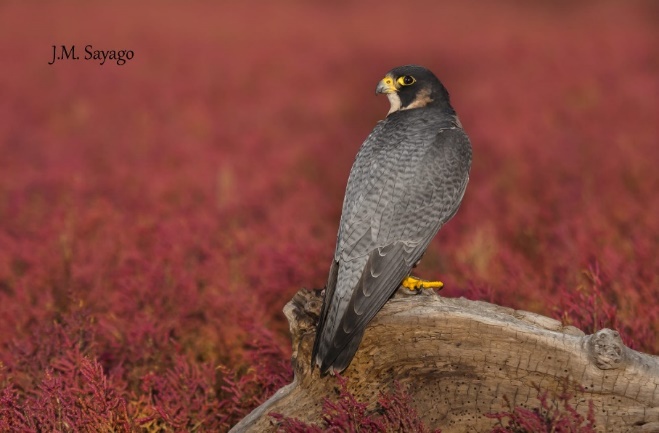

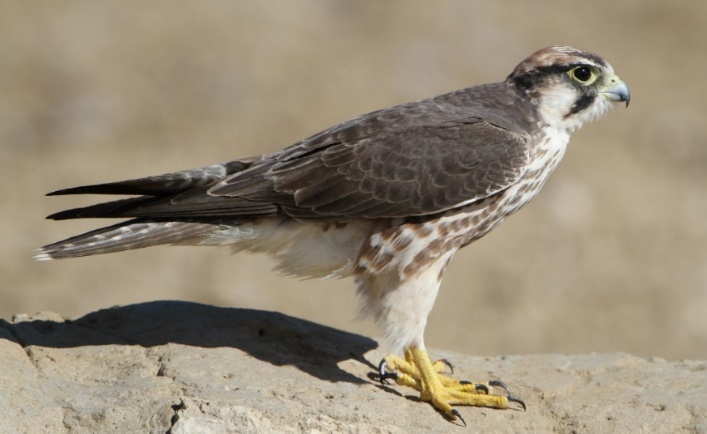

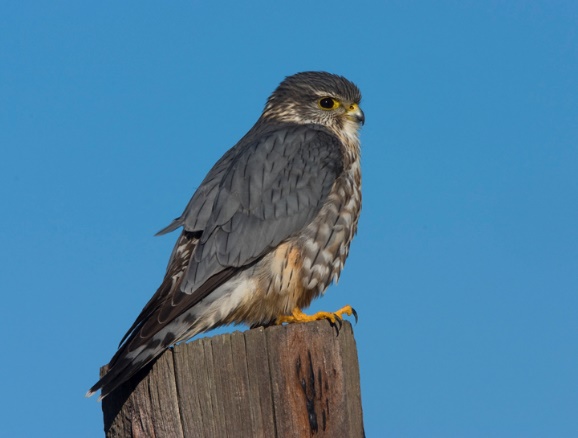
9

**Group 13**: *Falco tinnunculus* (left) and *F. naumanni* (center) by J.J. Negro. *F. vespertinus* (right) by B. Rodríguez.

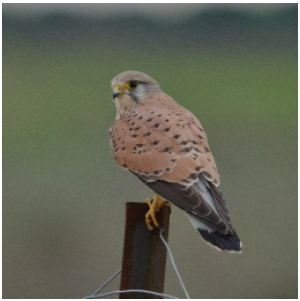

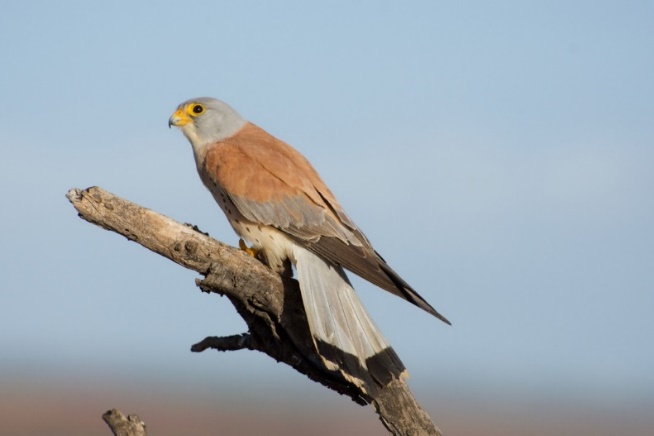

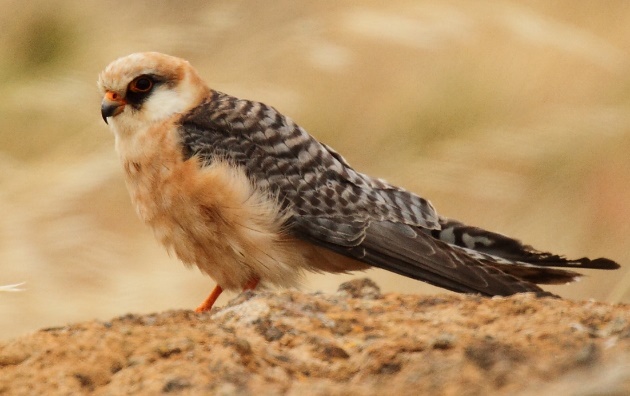

**Figure S2.** European raptors grouped according to trophic niche and phylogenetic relationships to show morphological and plumage similarities.

**Creative Commons attributions**

*-Haliaeetus albicilla* cropped from Christoph Müller (<https://commons.wikimedia.org/wiki/User:C-M>) under CC BY 4.0 (<https://creativecommons.org/licenses/by/4.0/legalcode>)

-*Accipiter brevipes* by Vladimir Yu. Arkhipov (https://commons.wikimedia.org/wiki/Special:Contributions/Arkhivov) under CC BY_SA 4.0 (https://creativecommons.org/licenses/by-sa/4.0/deed.en)

-*Buteo lagopus* by Marek Szczepanek (https://commons.wikimedia.org/wiki/User:Pkuczynski/Marek_Szczepanek) under CC BY_SA 3.0 (<https://creativecommons.org/licenses/by-sa/3.0/legalcode>)

-*Buteo rufinus* by Koshy Koshy (https://www.flickr.com/people/97235261@N00) under CC BY 2.0 (<https://creativecommons.org/licenses/by/2.0/legalcode>)

-*Clanga pomarina* by Wolbrun Avner (Creative Commons Attribution-ShareAlike 4.0 International)

-*Falco rusticolus* by NorthernLight (<https://de.wikipedia.org/wiki/Benutzer:NorthernLight>) under CC BY_SA 3.0 (<https://creativecommons.org/licenses/by-sa/3.0/legalcode>)

-*Falco biarmicus* by Derek Keats (https://www.flickr.com/people/93242958@N00) (https://es.wikipedia.org/wiki/Falco_biarmicus#/media/Archivo:Lanner_falcon,_Falco_biarmicus,_at_Kgalagadi_Transfrontier_Park,_Northern_Cape,_South_Africa_(34447024731).jpg) under CC BY 2.0 (<https://creativecommons.org/licenses/by/2.0/legalcode>)

-*Falco columbarius* by Gregory Smith (https://www.flickr.com/photos/22170893@N06/2276417896). [Creative Commons](https://en.wikipedia.org/wiki/en:Creative_Commons) [Attribution-Share Alike 2.0 Generic](https://creativecommons.org/licenses/by-sa/2.0/deed.en) license
